# High-speed, multicolor, structured illumination microscopy using a hexagonal single mode fiber array

**DOI:** 10.1101/2020.12.01.406652

**Authors:** Taylor A. Hinsdale, Sjoerd Stallinga, Bernd Rieger

## Abstract

Structured Illumination Microscopy (SIM) is a widely used imaging technique that doubles the effective resolution of widefield microscopes. Most current implementations rely on diffractive elements, either gratings or programmable devices, to generate structured light patterns in the sample. These can be limited by spectral efficiency, speed, or both. Here we introduce the concept of fiber SIM which allows for camera frame rate limited pattern generation and manipulation over a broad wavelength range. Illumination patterns are generated by coupling laser beams into radially opposite pairs of fibers in a hexagonal single mode fiber array where the exit beams are relayed to the microscope objective’s back focal plane. The phase stepping and rotation of the illumination patterns are controlled by fast electro-optic devices. We achieved a rate of 111 SIM frames per second and imaged with excitation patterns generated by both 488 nm and 532 nm lasers.

## Introduction

Understanding biology at the microscopic scale with chemical specificity is one of the *de-facto* driving forces behind fluorescence microscopy (1, 2). It has been an essential tool that has allowed the investigation of fundamental biological principles; however, the diffractive nature of light limits the resolution of classical light microscopy to ∼250 nm. Unfortunately, many interesting biological structures, such as the arrangement of the nuclear pore complex, lie beyond this limit. Imaging modalities such as electron microscopy can be used to study structures at these length scales, but they lack chemical specificity, and with it, the ability to infer information about their function.

A handful of techniques were introduced that fundamentally changed how the sub-diffractive arrangement and function of biological structures were studied (3–8). They are generally referred to as “super resolution microscopy” methods. The method of choice is dictated by the length and time scale of the phenomena being interrogated. For example, localization microscopy focuses on static biological constructs or single molecule tracking where resolution on the order of 10-20 nm is necessary. The fundamental limitation of localization microscopy is a sparsity constraint that drastically increases the time required to generate a reconstructed image, often on the order of 1-10 minutes, sometimes taking multiple hours(9). When imaging dynamic biological phenomena that occur throughout the cell, this is insufficient. In this regard, structured illumination microscopy (SIM) remains the dominant imaging method for dynamic cellular imaging at a resolution of ∼100 nm.

SIM was first introduced by *Heintzmann et al*. and *Gustafsson et al*. in 2000 (7, 10). The principles of SIM can be best understood in terms of signal processing and frequency aliasing. The widefield resolution of a microscope is defined by its point spread function, which is determined by its numerical aperture, *NA*, and the wavelength, *λ*, of the emitted fluorescence, as follows:

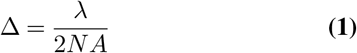

The Fourier transform of the point spread function, i.e. the optical transfer function (OTF), limits the band of detectable spatial frequencies in the image, where the smallest resolvable distance, Δ, corresponds to a spatial frequency of *k*_*max*_ = 1*/*Δ. This is commonly known as the Abbe diffraction limit. In 2D SIM, a well-defined fringe pattern, usually generated by two interfering plane waves, is projected into the sample via the excitation path of the microscope and then subsequently imaged by a pixelated detector. The superposition of the excitation pattern on the sample generates beat frequencies in the image which are known as Moiré fringes (7, 11). Spatial frequencies that reside outside of the OTF see the illumination pattern and are aliased into the pass band as beat, or difference, frequencies. Simply shown, the system can detect frequencies subject to |*k*_*ex*_−*k*_*s*_| ≤ *k*_*max*_, where *k*_*ex*_ is the spatial frequency of the excitation pattern, *k*_*s*_ is the spatial frequencies of the sample, and *k*_*max*_ is the maximum frequency of the system pass band. It then follows that the improvement factor goes as 2*k*_*ex*_*/k*_*max*_. The lateral resolution improvement can be extended to the third dimension by introducing axial modulation. This is typically done by interfering a third axially propagating beam with the two plane waves used in 2D SIM (12). Both 2D and 3D SIM have been widely used to study biological phenomena and structures *in vitro*. A broad overview of SIM’s biological applications can be found in(13–15).

Current SIM imaging systems utilize diffractive optical elements to generate structured illumination patterns in the sample. Although sufficient for static or slowly moving samples, using gratings to generate standing wave patterns is slow due to the need to physically rotate and shift the diffractive elements. The frame rate of such systems is often limited to 1 Hz for a full SIM acquisition (13). These systems can be made faster by using digitally programmable gratings, such as spatial light modulators (SLM) or digital micromirror devices (DMD); however, their multicolor utility is still limited by the highly wavelength dependent nature of diffractive elements. Different wavelengths will diffract off gratings at different angles, making it impractical to optimize a SIM system over a broad excitation wavelength range. This results in some wavelengths generating structured illumination patterns with larger than optimal pitches, which results in worse resolution improvement.

In this paper, we describe a SIM system that could be used for imaging dynamic media on the millisecond time scale with multiple excitation wavelengths. Our design utilizes a hexagonal array of single mode fibers to generate structured illumination patterns. The use of the hexagonal fiber array means that every wavelength can be optimized to generate its minimum pattern pitch in the sample, ensuring that reaching the maximum improvement factor for every excitation wavelength is easier. The speed of our SIM system is dictated by the pattern manipulation. The phase shifting and pattern switching are accomplished by fiber phase shifters and Pockels cells that operate well above 10 kHz, with a variable liquid crystal retarder limiting the polarization rotation to 2 kHz. To maximize the imaging speed of our system, we chose to first pursue 2D SIM over 3D SIM, an approach taken by other groups focused on high-speed imaging (16). This reduces the minimum number of images required for a single reconstruction from 15 for 3D SIM to 9 for 2D SIM. Although 3D SIM is often preferred over 2D SIM, especially in regard to its superior performance in thick samples, fast 2D SIM with millisecond time resolution can still be quite useful to researchers who are interested in dynamic imaging. We were able to image at well over 100 SIM frames per second and, with sufficient signal, it should be possible to exceed this.

## Materials and methods

### A. Optical system

The concept of our fast fiber SIM system is shown in Fig. 1. The optical system was constructed as an inverted microscopy platform to allow for easy imaging of mounted cells and other biological specimens. The illumination path consists of two single longitudinal mode lasers with differing wavelengths, one 488 nm (Sapphire SF NX 488, Coherent), and the other 532 nm (LCX-532S, Oxxius). The lasers are sent through an AOTF (AOTF 3151-01, Gooch & Housego) to enable high-speed wavelength switching and modulation. The lasers are then coupled into a polarization maintaining single mode fiber (PM-S405-XP, ThorLabs) via an achromatic fiber coupler (PAF2-A7A, ThorLabs). The fiber output is collimated using an achromatic lens (AC050-008-A, ThorLabs) and sent through a 50:50 non-polarizing beamsplitter (BS010, ThorLabs). The two beams emerging from the beamsplitter form the basis for generating a standing wave interference pattern in the sample. First, each beam is coupled into its own fiber phase shifter (FPS-001, General Photonics) to control the relative phase between them at up to a 20 kHz rate. In this way, the phase of the interference pattern can be controlled in the sample plane. The orientation of the interference pattern is controlled by coupling the two laser beams into differing pairs of fibers in a fiber array (SQS Vláknová optika a.s). The fiber array consists of 6 polarization maintaining fibers (PM-S405-XP, ThorLabs) arranged in a hexagonal pattern with each fiber at a vertex. The pattern is designed to directly replicate the Fourier plane of a single spatial frequency, standing wave pattern, arranged in three 120° spaced rotations. To create a standing wave, the light from the laser is coupled into two array fibers that are radially opposite one another, see Fig. 1. To switch between different pairs of fibers, thus controlling the pattern orientation, the two beams are sent through a large aperture Pockels cell (EM512, Leysop) and an achromatic quarter wave plate (AQWP05M-600, ThorLabs). This allows for greater than 10 kHz switching between S and P polarizations. A polarizing beam splitter (CCM1-PBS251/M,ThorLabs) is then used to switch the beams between two paths depending on the polarization state. Another Pockels cell and quarter waveplate are placed in series after the first pair, allowing for switching between three distinct beam paths. Each path contains two fiber couplers (PAF2-A7A, ThorLabs) and the appropriate fiber pair from the array. These fiber pairs are labeled as 1_a,b_, 2_a,b_, and 3_a,b_ in Fig. 1(a) and (b). The fiber array is then collimated using a 50 mm achromatic lens (ACA254-050-A, ThorLabs). All fibers in the array are oriented with the slow axes parallel to one another. To maintain a good pattern modulation contrast in the sample plane, the output polarization of the fibers needs to be rotated depending on the pattern orientation such that all light is S polarized in the sample plane. This is accomplished by using a polarization rotation unit which consists of a broadband wire grid polarizer (WP25M-VIS, ThorLabs), a high-speed liquid crystal retarder (HS LCVR, Meadowlark), and an achromatic quarter waveplate (AQWP10M-580, ThorLabs). This rotation unit is capable of providing halfwave rotation at greater than 1 kHz. The excitation beam is then relayed using two 180 mm achromatic lenses (AC508-180-A-ML, ThorLabs) to the first dichroic mirror (Di03-R405/488/532/635-t3-25×36, Semrock), which is used to compensate for phase lag between the S and P components of the excitation light generated by the main imaging dichroic (Di03-R405/488/532/635-t3-25×36, Semrock) (17). A third 180 mm achromatic lens then reimages the fiber array in the back focal plane of the 100X objective (CFI Apochromat TIRF 100XC Oil, Nikon). The +/-1 orders of the imaged array are focused to ∼90% of the spatial cutoff frequency of the system. This allows for the highest possible modulation pitch and the best resolution improvement for each color, while still being detectable in the raw images for pattern parameter estimation (12). The focused spots of the imaged fiber array in the objective back focal plane then become counter propagating beams in the objective’s sample focal plane. The emission is collected through the same objective in epi-illumination and the pupil plane is relayed to a deformable mirror (Multi-3.5, Boston Micromachines Corporation) via a tube lens (ITL200, ThorLabs) and a 125 mm achromatic lens (AC254-125-A-ML, ThorLabs) 4f pair. The emission path is then finally imaged onto an sCMOS camera (ORCA-Flash4.0 V2, Hamamatsu) by another 125 mm achromatic lens. All hardware is synchronized via a homebrew LabView application and two multichannel DAQ cards (M-Series, National Instruments).

**Fig. 1.**
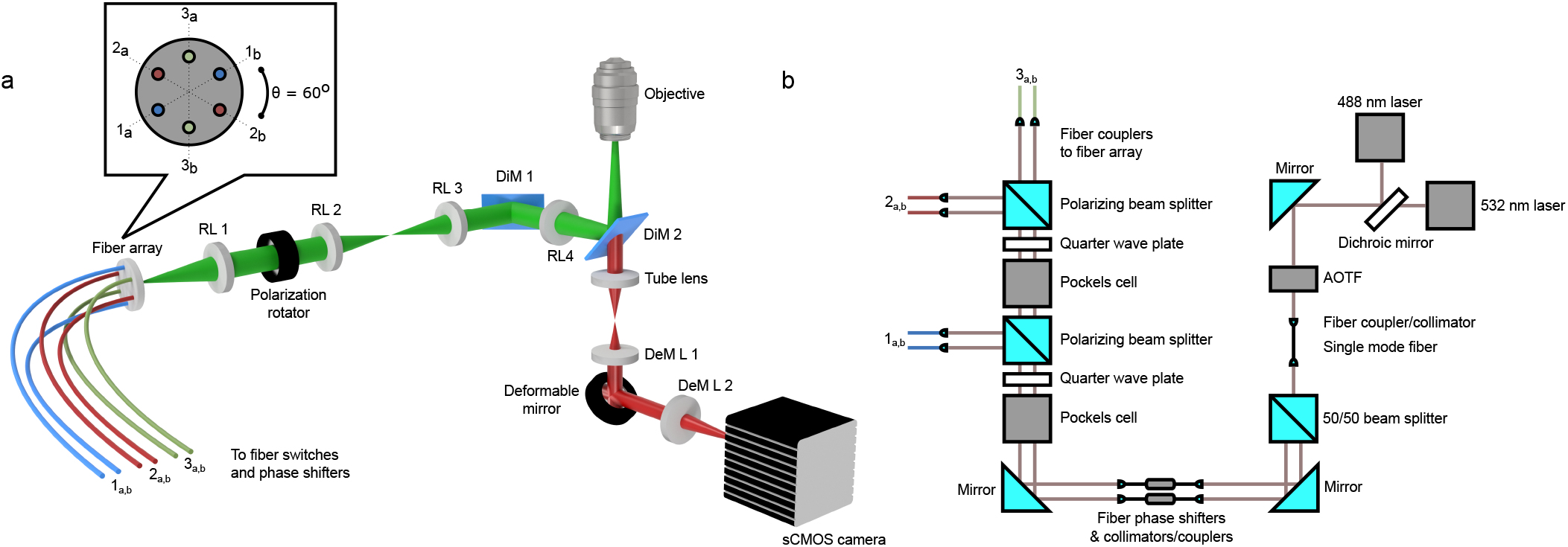
Fast fiber structured illumination concept. (a) shows a 3D rendering of the inverted microscope construction. The fiber array is collimated by relay lens RL 1 and then rotated by a polarization rotator. Lens pair, R 2&3, then relays the excitation beam to the compensation dichroic mirror (DiM 1). (RL 4) then focuses the light to the back focal plane of the objective after reflecting off of the imaging dichroic (DiM 2). The orientation of the pattern is determined by which pairs of fibers (1_a,b_, 2_a,b_, or 3_a,b_) are emitting light. The Fourier plane of the emission light is relayed by the tube lens and first deformable mirror lens (DeM L 1) to the deformable mirror. The abberation corrected emission light is then imaged onto the sCMOS camera via DeM L 2. (b) shows how the excitation lasers are phase shifted and switched between different fiber pairs.

### Fluorescent bead calibration samples

Fluorescent beads with a 100 nm diameter (F8800, ThermoFisher) were used to calibrate the optical performance of the microscope, asses the quality of the modulation pattern generated by the two interfering beams, and as a standard to evaluate the SIM improvement under ideal conditions. The protocols laid out in ref. (18) were closely followed to make calibration samples appropriate for SIM. Sparse, well separated bead samples, mounted in index matching oil, were used to assess the OTF of the system. This was done by taking through focus z-stacks with well-separated beads and fitting them with a vector point spread function model to quantify the system aberrations (19). Once quantified, the inverse aberrations were applied to the deformable mirror and another z-stack was made to confirm the improved image quality. The same sparsely populated bead samples were used to assess the modulation contrast of the SIM pattern. First, the phase-voltage responses of the fiber phase shifters were calibrated by linearly ramping the control voltage through multiple phase cycles. This allowed us to directly correlate the spatial phase shift in the sample plane to the control voltage. This process was carried out separately for each fiber phase shifter due to their slightly different phase-voltage responses. The modulation contrast of the excitation pattern can also be extracted from the fits used to find the phase-voltage relationship. Monolayers of 100 nm beads were used to quantify the performance of the fiber SIM microscope under ideal conditions. These monolayers were generated by diluting the stock concentrations of beads with ethanol in ratios of 1:5-1:10. Pipetting these solutions onto a cleaned coverslip creates regions of densely packed beads that are 1 layer in thickness as the ethanol evaporates. Near the edges of the uniform bead patches, the density becomes low enough to visualize individual beads. Both the whole field of view (FOV) and individual point sources can be visualized using these samples, making them ideal for SIM system calibration.

### In-vitro cell samples

Fixed and stained Cos-7 cells (GATTA-Cells 4C, GattaQuant) were used to represent realistic imaging conditions. The cells were stained for four structures (Nucleus: DAPI, Mitochondria: Anti-Tom20 with Alexa Fluor 488, Microtubules: Anti-Tubulin with TMR, Actin: SiR) and mounted in ProLong Diamond. Our microscope was able to image both the mitochondria and microtubule networks with the lasers available in our current setup.

### Escherichia coli sample preparation

E.coli cells with HU-mYPet labelled chromosomes (FW2179, derivative of E. coli K12 strain described previously(20)) were grown in liquid M9 minimal medium (Fluka Analytical) supplemented with 2 mM MgSO4, 0.1mM CaCl2, 0.4% glycerol (Sigma-Aldrich), and 0.1% protein hydrolysate amicase (PHA) (Fluka Analytical) overnight at 30°C. On the day of the experiment, the overnight culture was refreshed (1:100 vol) for 2.5 hours on fresh M9 medium at 30°C. Then 2ml of the culture was centrifuged at 6’000 rpm for 2min and concentrated 10 times to obtain a dense cell solution. For imaging, 1 *µ*l of the sample culture was pipetted onto a 35mm glass bottom imaging dish (Mattek) and immediately covered with a flat agarose pad, containing the above composition of M9 medium as well as 3% agarose. The cover was then sealed with parafilm to prevent evaporation.

### Reconstruction algorithm

We used the open source algorithm FairSIM (21). The SIM reconstructions shown in this paper were generated using the FairSIM ImageJ plugin. All reconstructions were made with a Wiener regularization parameter, *ω*, of 0.05 unless stated otherwise.

## Results

### SIM bead monolayers

A simple way to test the initial performance of our SIM system was to image a monolayer of sub-diffractive 100 nm beads. Figure 2(a) shows a 33×33 *µ*m, 512×512 pixel, image of a bead monolayer that was excited by a 532 nm laser. The raw images were taken with 20 ms exposures, which is equivalent to 5.5 SIM frames per second. The FOV is split between the SIM reconstruction (bottom right) and the the pseudo widefield reconstruction (top left), which was obtained by averaging the 9 raw frames. It clearly shows the improved resolution of the fiber SIM system when compared to diffraction limited imaging. Figure 2(b) is the Region Of Interest (ROI) highlighted in yellow from Figure 2(a), which is again split between the SIM reconstruction and the widefield reconstruction. Figure 2(c) quantifies the average resolution improvement over the whole FOV by computing the Fourier ring correlation (FRC) on two subsequent acquisitions of the scene shown in Fig. 2(a) (22). Figure 2(d) quantifies the improvement by looking at the Gaussian profile of an individual bead. The measured Full Width Half Maximum (FHWM) of the SIM and widefield reconstructions from Fig. 2(d) are 106 nm and 222 nm, respectively. The accompanying FRC resolution is calculated to be 111 nm for SIM and 235 nm for widefield, roughly agreeing with the reported FWHM improvement. One thing to note is that the fixed pattern noise of the sCMOS camera used in this experiment generates correlation between subsequent frames that is not related to the image content. This introduces some offset in the FRC which is seen in Fig. 2(c). The absolute resolution values reported in this case should be taken with some caution. Instead, the FRC is used to emphasize the average resolution improvement over the whole FOV.

**Fig. 2.**
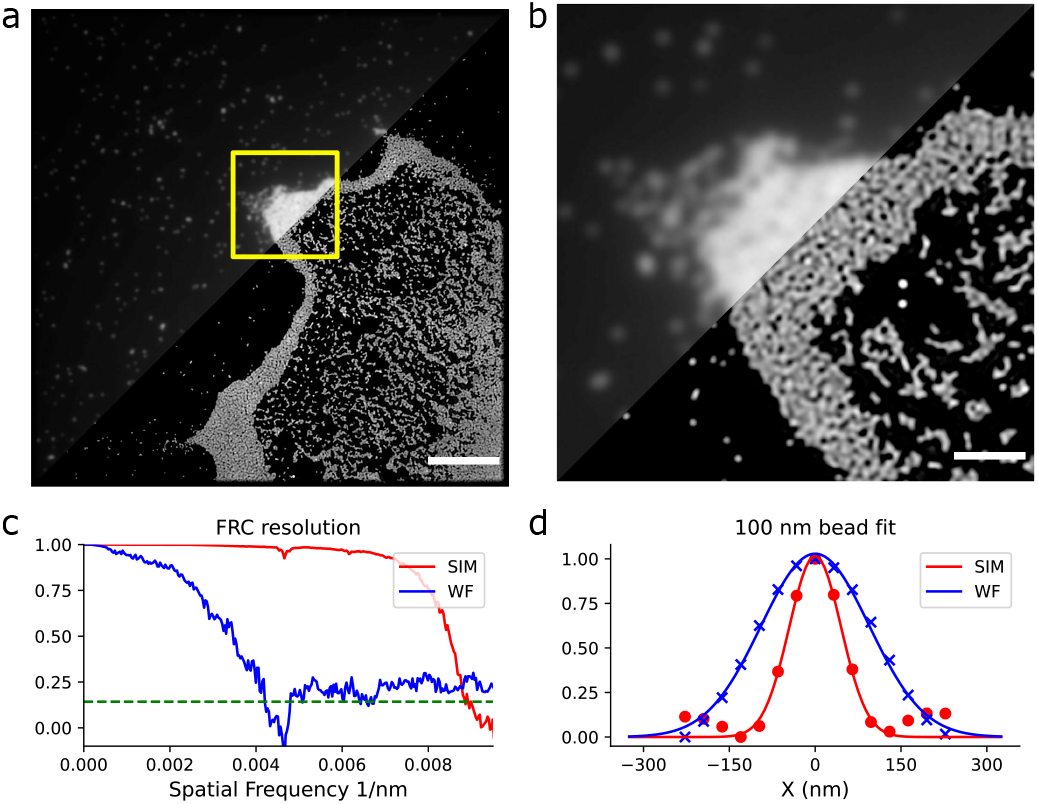
Assessment of quality of fiber SIM with 100 nm fluorescent bead sample. (a) 33×33 µm FOV with a monolayer of 100 nm beads, the scale bar is 5 µm. (b) This shows a zoomed in patch, denoted by the yellow square in a, the scale bar is 1.25 µm. The resolution improvement can clearly be seen in the SIM image where many individual, or aggregates of beads, are now distinguishable. (c) shows the FRC resolution for both the SIM and widefield images in (a). (d) shows the Gaussian fit to a single isolated bead for both the SIM and widefield reconstructions in (a)

The monolayers also provided an ideal sample to test the frame rate capabilities of the system. Figure 3(a) shows another typical 33×33 *µ*m monolayer FOV that is split between the SIM and widefield reconstructions. This was taken under 532 nm exciation with a 5 ms exposure per raw frame, for a total of 22 SIM frames per second. For many cellular imaging applications, this speed is sufficient. To test the limits of our system the exposure time was reduced. It should be noted that to increase the frame rate on an sCMOS camera, the FOV needs to be reduced to decrease the total pixel row readout time. Figure 3(b) shows an 8×8 *µ*m subsection from Figure 3(a) highlighted in yellow. Here, the raw frame exposure was set to 1 ms, for a total of 9 ms per SIM frame, amounting to 111 SIM frames per second, showing a 5-fold increase in SIM frame rate over the rate in Fig. 3(a). The 111 SIM frames per second movie can be viewed in Visualization 1. Figure 3(b) also shows an image series where the sample was laterally translated over a distance of 750 nm within 30 ms, implying a speed of *v* = 25 nm/ms. There are only 2-3 degraded reconstructions between the start and stop of the step, which is roughly 18-27 ms of lost data. The temporal shift is quantified in figure 2(c) where the displacement of the raw 1 ms frames is plotted. A smoothing spline was fit to the raw data and the settling time of the step response was calculated to be 20 ms. Figure 3(d) shows the FRC step response for the SIM reconstructions. This was done by taking the FRC between the *n* and *n* +1 frames in the reconstruction series. We estimate that the width of the FRC step response time series would be on the order of *λ/*(*NA*∗*v*)∼15 ms. A Gaussian function was fit to the time series and the 1/e width was found to be 23 ms. The speed of this FOV shift is larger than typical dynamic phenomena studied in SIM, such as cytoskeletal rearrangement (23). This opens up the opportunity to push SIM imaging into the domain of diffusion and other fast processes.

**Fig. 3.**
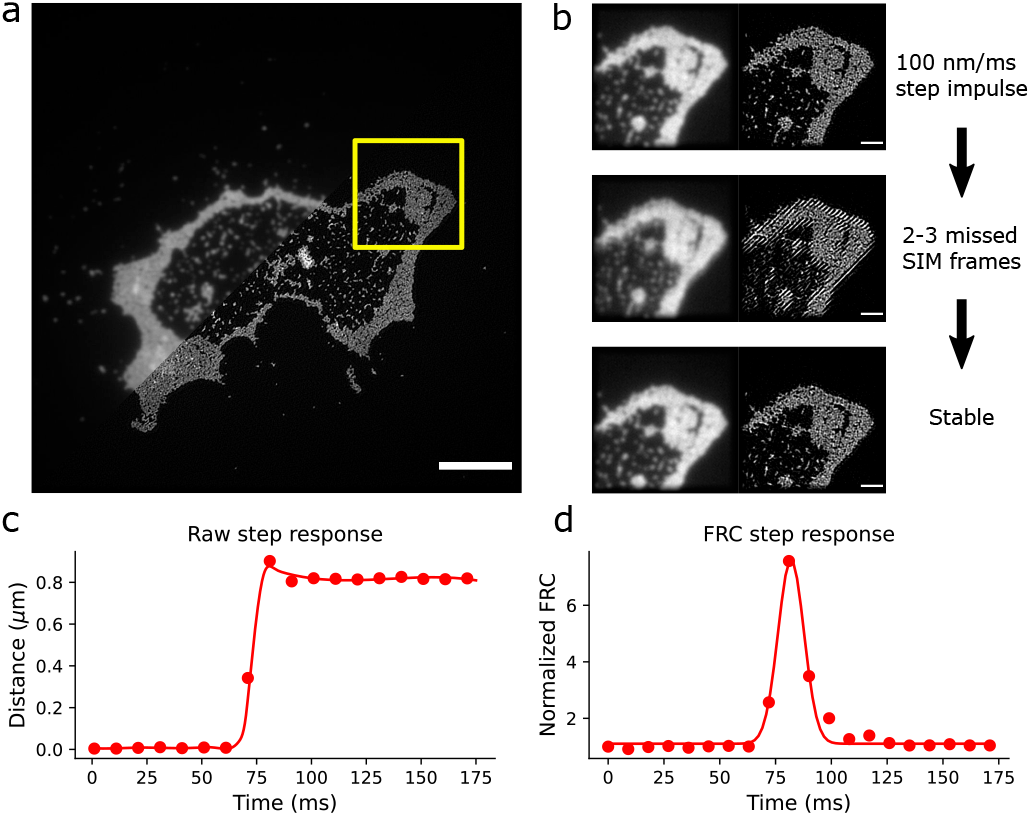
Assessment of the imaging speed of fiber SIM by rapid stage translation of 100 nm fluorescent bead sample. (a) 33×33 µm FOV with a monolayer of 100 nm beads, scale bar = 5 µm. (b) shows a zoomed in patch, denoted by the yellow square in a, scale bar = 1.25 µm. The resolution improvement can clearly be seen in the SIM image where many individual, or aggregates of beads, are now distinguishable. (c) shows the temporal response of the 1 ms raw image frames for the roughly 750 nm step. (d) plots the FRC as a function of time during the 750 nm step. The values are normalized by the static frame FRC value of 129 nm.

The absolute and relative improvements of the resolutions were also compared between Fig. 3(a) and (b) to show that the degraded Signal to Noise Ratio (SNR), due to lower photon counts, had a minimal effect on resolution improvement. The FRC resolution of Fig. 3(a) is 116 nm for the SIM reconstruction and 215 nm for the widefield reconstruction. The FRC resolutions for the 111 frames per second reconstructions in Fig. 3(b) are 111 nm for SIM and 218 nm for widefield. This shows that the absolute resolution remained similar as did the relative improvements. Similarly, the behaviours of the FWHM measurements in Fig. 3 are the same. The FWHMs of the SIM and widefield reconstructions from Fig. 3(a) are 96 nm and 215 nm, respectively. The FWHMs of Fig. 3(b) are 101 nm for SIM and 220 nm for widefield.

### Fixed in vitro COS7 cells

SIM has been extensively used to study fixed cells at sub-diffractive resolutions (15). We have investigated the fiber SIM system’s ability to super resolve both microtubules and mitochondria. Figure 4(a) shows another 33×33 *µ*m FOV of microtubules under 532 nm excitation, that is split between the widefield and SIM reconstructions. The raw frame exposure time was 40 ms, for a SIM frame rate of ∼3 frames per second. Figure 4(b) compares the smaller ROI highlighted in yellow from Figure 4(a). It is clear that the microtubules are better resolved, although their internal diameter of ∼25 nm is still too small for SIM. Figure 4(c) is a line plot of the profiles shown in Fig. 4(b). The tightly packed bundles of microtubules are largely blurred together in the widefield reconstruction, but almost every individual microtubule is resolved in SIM. Figure 4(d) shows a multicolor rendering of microtubules and mitochondria, where each color channel was acquired serially. The red color channel corresponds to microtubules under 532 nm excitation and the green channel corresponds to mitochondria under 488 nm excitation. The 33×33 *µ*m FOV here was significantly more dense than the pure microtubule FOV shown in Fig. 4(a). Because of this, the OTF attenuation option in FairSIM was utilized to reduce artifacts due to outof-focus fluorescence. The Wiener parameter here was left untouched, but the OTF attenuation settings for FairSIM were set so that the strength, *a*, was 1.0, and the width, FWHM, was 1.2. The exposure time for the raw frames was 40 ms, translating to ∼3 SIM frames per second. Figure 4(e) displays an enlarged ROI containing mitochondria that are outlined in yellow in Fig. 4(d). Two line profiles are drawn through one of the mitochondria in the ROI and plotted in Fig. 4(f), clearly indicating an improvement in resolution. The imaging speed limit in the cellular samples was also tested by reducing the FOV to 16×16 *µ*m (256×256 pixels) and decreasing the exposure time to 2 ms. This gave an effective SIM frame rate of 55 frames per second. Figure 4(h) shows the results of this on a region of microtubules. Here one can see the reconstructed widefield image on the left and the SIM reconstruction on the right with no qualitative degradation when compared to the longer exposed images of Figs. 4(a) and (d). Figure 4(i) quantifies the resolution using FRC analysis. The resolution is comparable to all other FRC resolution measurements at this excitation wavelength, 532 nm, with SIM having a FRC value of 119 nm and the widefield having a value of 219 nm.

**Fig. 4.**
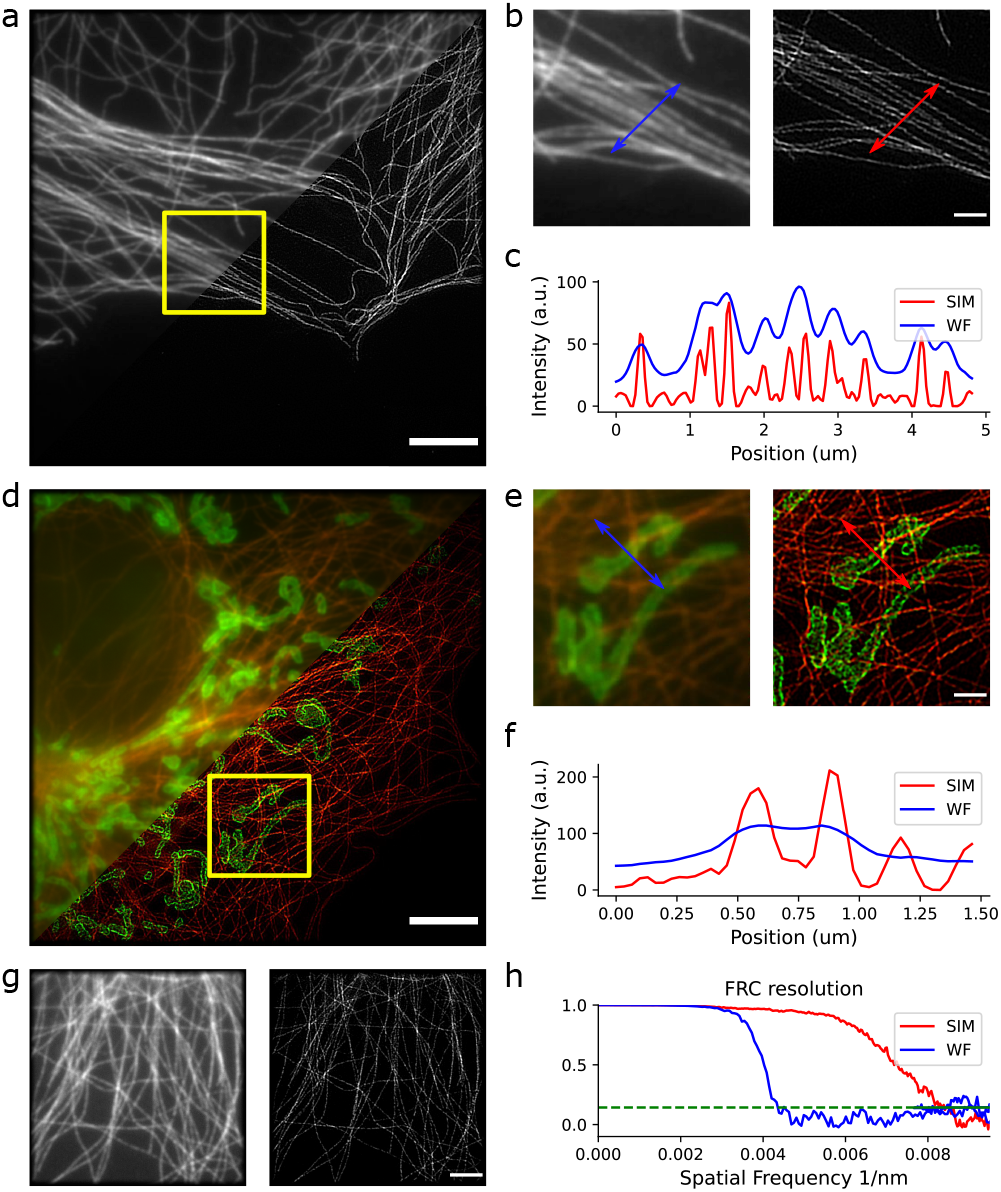
Fiber SIM multi-color imaging of fixed COS7 cells. (a) shows a 33×33 µm FOV image of a microtubule network. Half of the image is displayed as the widefield reconstruction and the other half as the high-resolution SIM reconstruction. The scale bar is 5 µm. (b) shows the region cropped by the yellow square in (a). Here it is easy to compare the apparent resolution improvement. The scale bar is 1.25 µm. (c) plots a line profile through the microtubules and clearly shows the resolution improvement. (d) show multi-color capabilities of the system. It is a 33×33 µm FOV with microtubules displayed in red and mitochondria displayed in green. The scale bar is 5 µm. (e) shows the region cropped by the yellow square in (d) with the scale bar equal to 1.25 µm. (f) plots a line profile through a mitochondria. (g) displays a high-speed acquisition, 55 SIM frames per second, of a microtubule network. The left panel shows the widefield reconstruction and the right panel shows the SIM reconstruction. The scale bar is 2.5 µm. (h) shows the FRC resolution of the highspeed images in (g) to confirm that the resolution improvement is preserved

### Live in vitro e. coli

The fiber SIM system was used to image live cellular samples to test it under poorer imaging conditions. These conditions were caused mostly by using genetically encoded fluorescent proteins, which are typically less bright than organic fluorophores, and the lack of an anti-fade mounting medium to scavenge oxygen and prevent photo-bleaching. Figure 5(a) shows a 33×33 *µ*m FOV of *e. coli* whose genome has been encoded to express yellow fluorescent protein. These images were taken with 40 ms exposures for the raw frames under 532 nm excitation. The OTF attenutation function in FairSIM was again used with *a* = 1.0 and the FWHM = 1.2. A series of 11 widefield and 11 SIM reconstructions were made and averaged to form the images in Fig. 5. The effect of this averaging is that the high-frequency reconstruction noise is reduced, and the common structure in each image series is preserved. Many of the cells appear to be uniform in intensity, but some show a structured and condensed chromosome. Figure 5(b) shows the yellow highlighted ROI from Figure 5(a). The left panel of Figure 5(b) is the average of 11 widefield reconstructions and the right panel is the average of 11 SIM reconstructions. Here one can see a “figure eight” shape, which may be a replicating chromosome (24). Figure 5(c) shows a line profile through a loop in the figure eight for both the averaged widefield and SIM images. In the widefield image, this is completely blurred together and most of the spatial information is lost. The SIM reconstruction clearly shows a separation. The confidence of this structure is further enhanced by comparing the average of the SIM reconstructions to the 11 individual SIM reconstructions in Visualization 2. As the noise in the underlying raw images of the different SIM reconstructions is independent then so is the typical SIM reconstruction noise in the resulting SIM reconstructions. It may therefore be concluded that the variability over different reconstructions is an objective measure for the SIM reconstruction noise. Blob-like features that are consistently present in all reconstructions may be considered trustworthy.

**Fig. 5.**
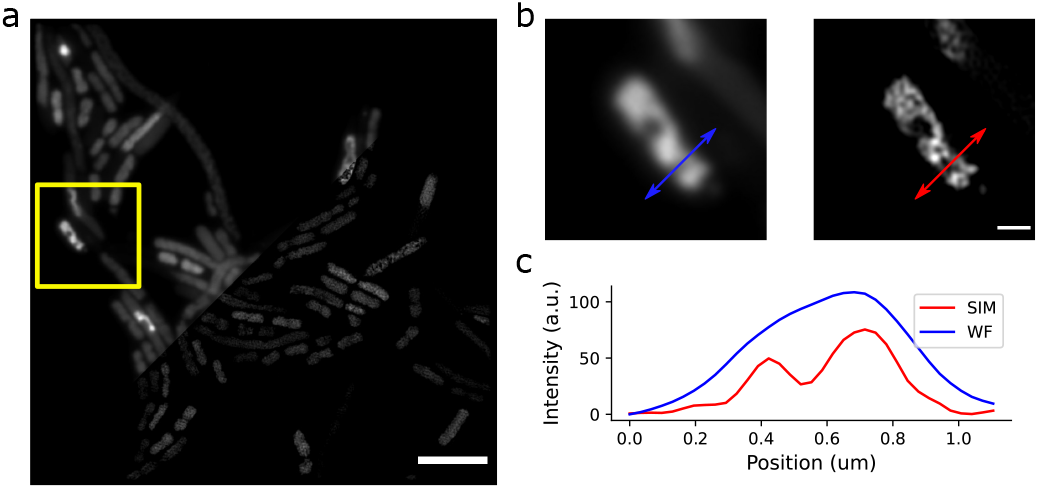
Fiber SIM imaging of live e. coli bacteria. (a) shows a 33×33 µm FOV image containing several bacteria. Half of the image is displayed as the widefield reconstruction and the other half as the high-resolution SIM reconstruction. The scale bar is 5 µm. (b) shows the region cropped by the yellow square in (a). This is hypothesized to be multiple entangled chromosomes. The scale bar is 1.25 µm. (c) plots a line profile through a loop in the chromosome.

### Fiber SIM speed

The system’s speed is directly related to how fast the sCMOS camera can be read out. The speed of the camera is in turn limited by the desired FOV and the exposure time. This also determines the effective exposure time, or global exposure, which is the difference between the set exposure time and and pixel row readout time of 9.65 *µ*s. The relationship can be written as *T*_*global*_ = *T*_*exposure*_−*T*_*read*_, where *T*_*read*_ = *H*_*lines*_*/*2. This has implications when trying to increase the speed while maintaining the same FOV. The period of global exposure will become so brief, that there will be very little illumination, thus tying the frame rate and FOV together even more closely. For the current setup, the illumination profile was designed to provide homogeneous illumination over a ∼30-50 *µ*m diameter, or 512×512 pixels. The camera limited speed of the fiber array system is summarized in Table 1, starting with a FOV of 512×512 pixels. Please note that this is the maximum achievable speed at a given FOV; the exposure time can always be made longer for slower acquisitions. Achieving the maximum speed at 64×64 pixels can also prove challenging as the response time of the liquid crystal rotator becomes a limiting factor.

**Table 1.** Camera limited fiber array SIM speed

| FOV (pixels) | FOV ($\mu\text{m}$) | Raw fps | SIM fps |
| --- | --- | --- | --- |
| 512x512 | 33x33 | 405 | 45 |
| 256x256 | 16x16 | 810 | 90 |
| 128x128 | 8x8 | 1620 | 180 |
| 64x64 | 4x4 | 3240 | 360 |

## Discussion

### Maximum realizable speed

The advantages of fiber array based structured illumination are presented in this paper. It was shown that fiber array SIM allows for high-speed pattern switching with electro-optics, and fast phase modulation with kilohertz fiber phase shifters. The result is that the structured illumination data acquisition becomes limited by a few key hardware elements. In our case, the main limiting factors were the liquid crystal polarization rotator (LCPR) and the frame rate of the camera. Many home-built SIM setups use LCPRs to corotate the polarization of the excitation beam with the SIM pattern rotation to ensure that the interference pattern is always S polarized in the sample plane (17, 23). These are typically limited to 1 kHz rotation speeds; however, some devices are able to push the limit with a 500 *µ*s switching speed, giving a ∼2 kHz rotation rate. There are a few options that can be explored to remove the necessity of one of these devices. For systems based on Spatial Light Modulators (SLMs) and Digital Micromirror Devices (DMDs), if only 2D SIM is being pursued, various groups have used a quarter waveplate to generate circularly polarized light after the diffractive device (25). The diffracted beams are then sent through a “pizza” polarizer, which is a multi-segmented polarizer cut into wedges (26, 27). Each wedge is aligned on a circle such that the polarization axis is perpendicular to the radius, creating an S polarized interference pattern at the sample. This is a simple solution for 2D SIM with the caveat that it throws away half of the available excitation light. An alternative would be to use a fiber array-based SIM system and prealign the fiber polarization axis in the same way the pizza polarizer is aligned. This would essentially eliminate all frame limiting factors besides the camera itself.

### Camera limitations

Current generation sCMOS cameras are largely similar in design and based on similar sensor arrays. Because of this, the limiting factor will always be the pixel line readout time, ∼9.65 *µ*s. Some manufacturers “sandwich” two sCMOS arrays together and read them from the middle out, effectively doubling the camera frame rate. Intensified high speed CMOS cameras could be considered as a viable alternative to sCMOS cameras for high speed, low light, imaging(28). Unfortunately, their relatively large pixel size would diminish the speed increase due to Nyquist sampling requirements. Another, more complex, option is to multiplex each phase and orientation onto different regions of interest on the camera (29, 30). This could potentially push SIM imaging speeds to over 10 kHz for smaller fields of view. Fiber array SIM would be extremely advantageous in this regime because the pattern switching and phase shifting can easily exceed 10 kHz. When compared to SLM based systems, even the fastest modern SLMs are limited to ∼4 kHz. DMD devices could be used instead of SLMS, but their low diffraction efficiency and highly wavelength dependent behaviour make them a non-ideal choice for widefield SIM with multiple colors.

### Comparison to state-of-the-art

Current state-of-the-art systems for multicolor SIM imaging at high speeds utilize SLMs(16). The work published by *Markwirth et al*. shows a binary ferro-electric SLM based SIM system that was nearly camera frame rate limited. The maximum reported frame rate for a single color channel was 57.8 SIM frames per second over a 256×256 pixel, 20*µ*mx20*µ*m, FOV. This was accomplished by imaging with 0.5 ms raw frame exposures at 520 frames per second. With this system, they were able to show freely diffusing fluorescent microspheres. A main hardware limitation of their system is the SLM device which has a maximum pattern refresh rate of 0.44 ms, or 2272 Hz. Our fiber-based SIM system compares favorably on this account. As discussed previously, our pattern switching and phase shifting is capable of exceeding 10 kHz due to its electro-optic construction; however, we would need to replace the LCPR in our system with the “pizza-polarizer” setup discussed in section A to fully take advantage of our fast pattern manipulation.

### 3D fiber array SIM

3D SIM is commonly used in cellular research for its superior background rejection and axial resolution enhancement when compared to 2D SIM. This is accomplished in commercial systems by either using a diffraction grating or an SLM, and interfering the +/-1 orders with the 0th order. This creates what is known as a “woodpile” pattern, which is an axially varying standing wave. This can be done by utilizing a fiber array with a central fiber to act as the 0th diffraction order. One crucial factor is that the precise phase relation between the +/-1 orders and the 0th order must be controlled. If this is not known, the standing wave pattern can be displaced axially and not reside at the optimal position for 3D SIM. In diffraction based systems, this problem is trivial because the diffractive element acts as a common path interferometer, ensuring that the phase relationship between every order is stable and known. For 3D fiber array SIM, the relative phase of all diffracted orders must be monitored and actively controlled. Another complication is that 3D SIM requires, at a minimum, 5 phases and 3 angles for a complete reconstruction. This lengthens the acquisition time beyond the 9 frames required for 2D SIM by ∼60%. Additionally, if volumetric z-stacks are required, the increased resolution in the axial direction raises the sampling requirements when creating focal stacks by a factor of two. All together, this roughly doubles the imaging time required. The challenges of implementing 3D fiber SIM also come with some unique opportunities not afforded to grating, DMD, and SLM based systems. Specifically, the axial modulation is no longer fixed relative to the objective focal plane. The phases of the three illuminating fibers could be individually controlled to allow for precise positioning of the “woodpile” pattern in 3D space, which would enable quick remote focusing to any arbitrary z-plane. This could be quite useful when imaging moving objects, such as bacteria, that quickly displace in all dimensions. The independent phase control would also allow for compensation of depth-dependent spherical aberration, ensuring an optimum modulation depth of the standing wave over a larger range of imaging depths(31).

## Conclusion

In this paper we have shown a novel implementation of structured illumination microscopy using a single mode fiber array to perform fast super resolution imaging in multiple colors. Fast manipulation of the pattern phase was achieved by utilizing inline fiber phase shifters and manipulation of the pattern orientation was achieved by using a series of Pockels cells, waveplates, and polarizing beam splitters. Our fiber SIM method demonstrated the ability to improve the resolution over standard widefield microscopy in fluorescent bead samples, fixed cellular samples, and live cellular samples. It was able to achieve fast imaging rates up to 111 SIM frames per second, thus approaching a camera frame rate limited system and pushing SIM towards new speed boundaries.

## Supporting information

Visualization 1

Visualization 2

## Acknowledgments

Live *e. coli* samples were provided by the lab of Cees Dekker at the Delft University of Technology. All *e. coli* sample preparation was performed by Aleksandre Japaridze and Milos Tisma.

